# Interplay of TGFb superfamily members governs optic fissure closure

**DOI:** 10.1101/010652

**Authors:** Stephan Heermann, Priska Eckert, Juan L. Mateo, Eleni Roussa, Belal Rahhal, Aimee Zuniga, Kerstin Krieglstein, Jochim Wittbrodt

**Affiliations:** Centre for Organismal Studies (COS) Heidelberg, Im Neuenheimer Feld 230. 69120 Heidelberg, Germany; Department of Molecular Embryology, Anatomy, Institute of Anatomy and Cell Biology, University Freiburg, Albertstrasse 17, 79104 Freiburg, Germany; Developmental Genetics, University of Basel Medical School, 4058 Basel Switzerland; present address: Anatomie und Zellbiologie, Heidelberg University INF 307, 69120 Heidelberg, Germany; present address: School of Medicine and health sciences, An-Najah National University, Nablus, Palestine

**Author notes:** Authors for correspondence: Stephan Heermann Joachim Wittbrodt.

## Abstract

The optic fissure is a gap in the developing vertebrate eye and must be closed as development proceeds. A persisting optic fissure is referred to as coloboma, a major cause for blindness in children. Multiple factors have been linked to coloboma formation, however, the actual process of fissure closure is only poorly understood.

Based on our findings we propose an important role of TGFb signaling for optic fissure closure. We show active TGFb signaling in the fissure margins, analyzed by a new TGFb signaling reporter zebrafish. We found BMP antagonists regulated by TGFb. These antagonists we also found expressed in the fissure margins. Finally we show a coloboma phenotype in a TGFb KO mouse. Microarray data analysis indicates intense TGFb dependent remodeling of the extracellular matrix (ECM) during optic fissure closure.

We propose that TGFb is driving optic fissure closure by ECM remodeling. As previously shown, inhibition of BMP signaling is important for such TGFb dependent ECM remodeling. We show that this is achieved by the regulation of BMP antagonists, expressed in the optic fissure margins.

## Introduction

Early vertebrate eye development starts already at late gastrula, by the definition of the eye filed in the prosencephalon. The evagination of the optic vesicles (ov) on both sides is based on single cell migration (Rembold et al., 2006). The early eye filed can already be identified by the expression of retinal homedomain transcription factor (rx) genes and loss of rx gene function results in anophtalmia (Loosli et al., 2003, Loosli et al., 2001).

The optic vesicle is surrounded by periocular mesenchyme (POM) and gets in close proximity to the surface ectoderm, where the lens is being induced. Subsequently the optic vesicle is being transformed into the optic cup (reviewed in Fuhrmann 2010), which is bending around the forming lens (Martinez Morales et al., 2009, Bogdanovic et al., 2012). Recently we observed that the lens-averted part of the optic vesicle is largely integrated into the forming neuroretina during this process (Heermann et al., 2014). In addition, we showed that the forming optic fissure at the ventral pole of the optic cup serves as an important gap for the lens-averted epithelium to flow into the optic cup to constitute the ventral part of the optic cup (Heermann et al., 2014). Notably this epithelial flow is dependent on the presence of a BMP antagonist (Heermann et al., 2014). BMP signaling and also BMP antagonism are known to be important for vertebrate eye development at different developmental stages (Morcillo et al., 2006, Koshiba-Takeuchi et al., 2000, Sasagawa et al., 2002, Behesti et al., 2006, Sakuta et al., 2001, Heermann et al., 2014).

BMP, expressed in the dorsal part of the optic cup, together with Vax2, expressed in the ventral part of the optic cup, were shown to define the dorso-ventral identity of retinal domains (Koshiba-Takeuchi et al., 2000, Sasagawa et al., 2002, Mui et al., 2005, Behesti et al., 2006).

Importantly the optic fissure at the ventral pole of the optic cup is only a transient structure and the fissure margins have to fuse as development proceeds to generate a continuous ventral retina for perception of the dorsal visual field. Failures in optic fissure closure can affect all tissues of the eye and are termed coloboma (reviewed Gregory-Evans et al., 2004, 2013).

Here we show BMP antagonists expressed along the optic fissure margins opposing the BMP ligand, which is expressed in the dorsal optic cup. We found the expression of the BMP antagonists to be dependent on TGFb signaling with TGFb ligands expressed in the developing lens and periocular tissue as well as their ligand binding receptor, TGFBR2 (TBRII), in the optic fissure region. With the help of a newly generated TGFb-signaling reporter zebrafish line we identified active TGFb signaling in the developing optic fissure margins, matching the expression of the BMP antagonists. To further aim for the functional implication of TGFb signaling in optic fissure fusion we analysed a TGFb KO mouse and found a failure in optic fissure fusion, a coloboma.

## Results

### BMP and BMP antagonism in the developing eye

The specification of dorsal versus ventral cell identities within the vertebrate eye is governed by BMP4 and Vax2 (Koshiba-Takeuchi et al., 2000, Sasagawa et al., 2002, Mui et al., 2005, Behesti et al., 2006). BMP4 is expressed in the dorsal optic cup (Fig 1A) whereas Vax2 is expressed in the opposing ventral optic cup (Fig 1B). BMP4 as a secreted ligand of the TGFb superfamily can potentially travel over long distances. Therefore it is an important finding that BMP antagonists, gremlin 2 (grem2) (Fig 1C) and follistatin a (FSTA) (Fig 1D), are also expressed in the ventral optic cup opposing the BMP4 ligand. Notably, the expression is further defined to the margins of the optic fissure (Fig 1 C/D). This very precisely defined expression of BMP antagonists at the optic fissure margins suggests that BMP signaling is actively inhibited at this site.

**Figure 1:**
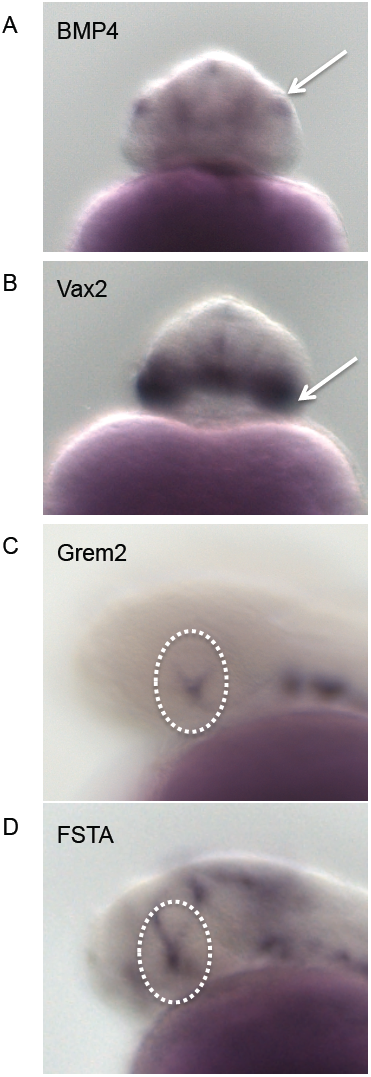
BMP antagonists are expressed in the optic fissure margins opposing the dorsally expressed BMP4 (**A**) in-situ hybridization of BMP4 (±30hpf zebrafish), frontal view. Note the localized dorsal expression of BMP4 within the optic cup (arrow). (**B**) in-situ hybridization of Vax2 (±30hpf, zebrafish), frontal view. Note the localized ventral expression of Vax2 within the optic cup (arrow). (**C**) in-situ hybridization of Grem2 (±30hpf, zebrafish), lateral view. Note expression of Grem2 in the optic fissure margins (encircled). (**D**) in-situ hybridizations of Fsta (±30hpf, zebrafish), lateral view. Note expression of Fsta in the optic fissure margins and the developing CMZ (encircled).

### Expression of TGFb pathway members during in the developing eye

Notably Zode and colleagues (Zode et al., 2009) described an interaction between TGFb and BMP in cells of the optic nerve head in the context of glaucoma. TGFb was found to induce Gremlin to inhibit BMP signaling. This in turn released the inhibition of BMP on TGFb induced extracellular matrix (ECM) remodeling. Since during development the optic nerve head originates from the most proximal domain of the optic fissure, we reasoned that similar regulatory circuits could also be controlling the closure of the optic fissure during development. Therefore we investigated the expression of TGFb ligands and the TGFb ligand binding receptor during this stage of eye development. We found TGFb2 expressed mainly in periocular tissue (Fig 2A) whereas TGFb3 was expressed mainly distally in the developing lens (Fig 2B). The ligand binding receptor TBRII is stable on the level of the protein and heavily recycled. Therefore it is likely that RNA for this receptor is only expressed at low levels. We observed a weak expression of the TBRII at the site of the optic fissure (Fig 2C). In the next step we were interested whether TGFb signaling was active in the optic fissure margins.

**Figure 2:**
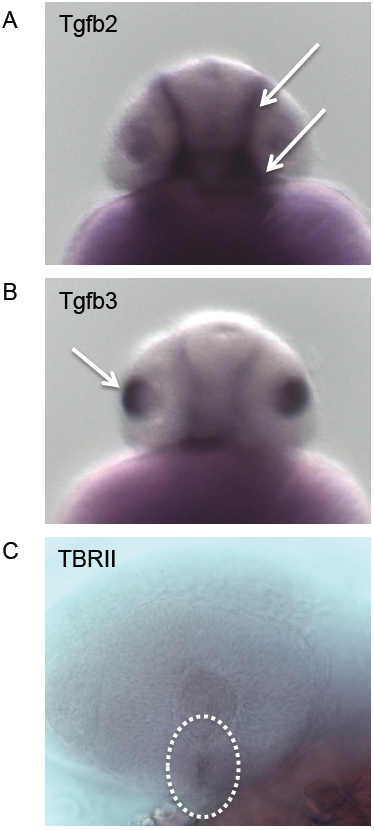
Expression of TGFb ligands and TGFb ligand binding receptor in the optic cup (**A**) in-situ hybridization of Tgfb2 (±30hpf, zebrafish), frontal view. Note expression of Tgfb2 in the periocular mesenchyme (arrows). (**B**) in-situ hybridization of Tgfb3 (±30hpf, zebrafish), frontal view. Note expression of Tgfb3 in the distal part of the developing lens (arrow). (**C**) in-situ hybridization of TBRII (±30hpf, zebrafish), lateral view. Note expression of TBRII in the optic fissure region (encircled).

### Zebrafish reporter line for TGFb signaling

In order to assess active TGFb signaling during zebrafish development we established a TGFb signaling reporter line. The reporter is based on the canonical transcription factors, Smads, by which TGFb signaling is transduced (Heldin et al., 1997). We used repetitive Smad Binding Elements (SBE) from the human plasminogen activator inhibitor (PAI). Such a reporter has been intensively used for years as a luciferase assay to assess the amount and activity of TGFb in cell culture (Dennler et al., 1998) and more recently in mice (Lin et al., 2005). We cloned the repetitive SBEs together with a minimal promoter element into an expression vector to drive the expression of a membrane bound GFP (GFPcaax) (Fig 3A) and generated a zebrafish line. Activated TGFb signaling can be observed during development, example given by domains in the forehead region as well as the distal tail (Fig 3B). To further test the functionality of this reporter line we wanted to see, whether signaling could be inhibited by a known TGFb signaling inhibitor, SB431542 (Inman et al., 2002, Laping et al., 2002). We observed a drastic reduction in signal activity after treatment with SB431542 (Fig 3C).

**Figure 3:**
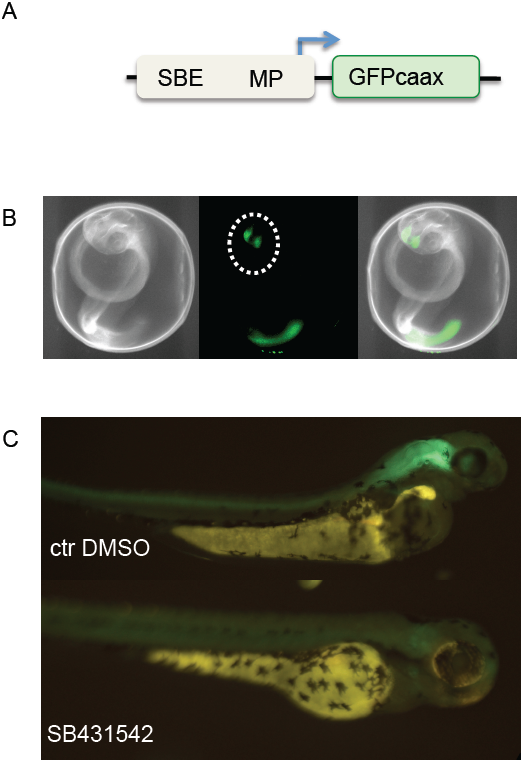
Establishment of a TGFb signaling reporter in zebrafish (**A**) construct generated for the TGFb signaling reporter. Multimerized Smad Binding Elements (SBEs) in combination with a minimal promoter (MP) drive membrane localized GFP (GFPcaax). (**B**) Brightfield, fluoreszent and merged images of a zebrafish larvae expressing the TGFb signaling reporter construct. Note the expression domains in the forebrain (encircled) and the tail. (**C**) In order to test the functionality de-chorionated zebrafish larvae expressing the TGFb signaling construct were exposed to a TGFb signaling inhibitor (SB431542). The inhibitor suppressed the activity of the TGFb signaling reporter dramatically (recorded at postembryonic stages). Please see the DMSO treated fish as control.

### Active TGFb signaling at the optic fissure margins

With the help of the generated TGFb signaling reporter zebrafish line we further analyzed the development of the optic fissure. In combination with a reporter line for shh signaling (Schwend et al., 2010) to identify the optic stalk we performed time lapse analyses and followed the signaling reporters during eye development (Fig 4 A-H). We found the TGFb signaling reporter first active in an area at the anterior-most tip of the embryo (Fig 4E). Subsequently TGFb signaling was activated in a region of the forebrain adjacent to the developing eye (Fig 4 B, C, F and G). Remarkably this region of activated TGFb signaling was eventually extending into the ventral optic cup, specifically into the forming optic fissure margins (Fig 4 D and H). This clearly indicates that TGFb signaling is active in the optic fissure margins.

**Figure 4:**
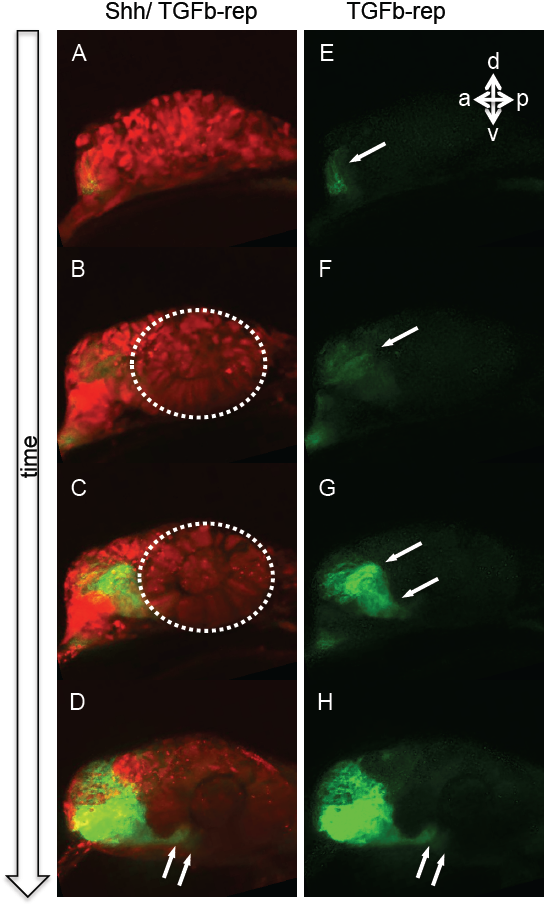
TGFb signaling is active in the developing optic fissure (**A-D**) and (**E-H**) progression of shh (red) and TGFb signaling (green) and TGFb signaling alone, respectively during optic cup (encircled in B-C) development. Note that the TGFb signaling activity spreads from the forebrain (arrow in E-G) into the developing optic fissure (arrows in D and H).

### Functional relevance of TGFb signaling for optic fissure closure

Having shown active TGFb signaling in the optic fissure margins we wondered what function TGFb exerts at this site. Since TGFb is important for ECM remodeling (Ignotz and Massague., 1986, Roberts et al., 1992, Zode et al., 2009, Sethi et al., 2011), we reasoned that TGFb could be important for the ECM remodeling during the fusion of the optic fissure. In mouse three TGFb ligands (TGFb1, 2 and 3) exist. In TGFb2 KO mice several phenotypes affecting the eye were reported (Sanford et al., 1997), examples given, a remaining primary vitreous, a peters anomaly like phenotype and a dyslayered neuroretina. Although a coloboma has not been reported, we analyzed eyes of TGFb2 KO embryos and specifically focused on the optic fissure region. Remarkably, in addition to the already described phenotypes we observed an affected closure of the optic fissure, a coloboma (Fig 5B, see A as control). Notably the optic fissure margins were able to get in close proximity to each other, but rather grew inwards than fusing orderly (Fig 5B). Furthermore we aimed at the expression levels of gremlin and follistatin in eyes of TGFb2 KO embryos. To assess these expression levels we analyzed the quantitative levels of RNAs prepared from E13.5 embryonic eyes by Agilent microarray chips. We compared TGFb2 KO (TGFb2/ GDNF double KO, Rahhal, Heermann et al., 2009) to wildtype embryos. We found gremlin1 (Fig 5C) as well as follistatin (Fig 5D) significantly reduced in eyes of TGFb2 KO mice. Taken together this strongly suggests that TGFb2 plays a crucial role during the closure of the optic fissure likely by the induction of BMP antagonists.

**Figure 5:**
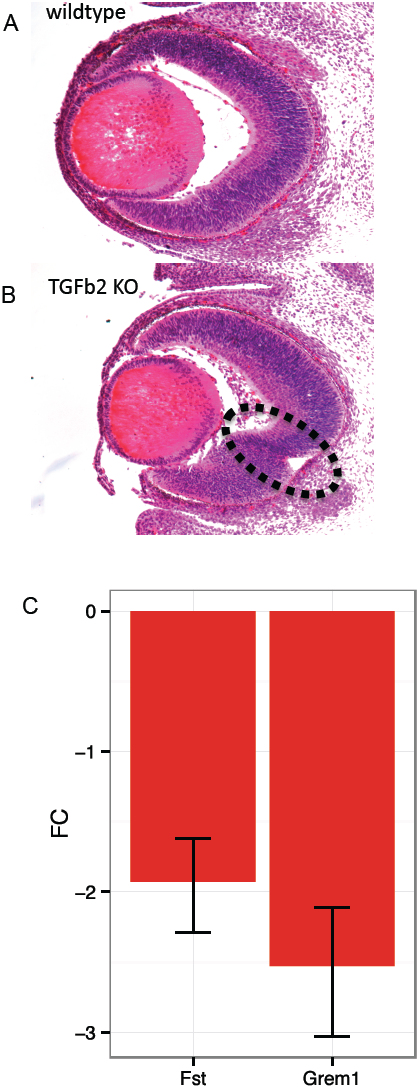
loss of TGF2 ligand results in coloboma (**A**) Frontal section of a wildtype embryo (E14.5) as control (H&E staining). (**B**) Frontal section of a TGFb KO embryo (E14.5) (H&E staining). Note the remaining optic fissure (encircled). Importantly the fissure margins must have met, but rather than fusing grew inwards. (**C**) Expression analysis of gremlin and follistatin, both being BMP antagonists, revealed a decrease in TGFb2 KO (TGFb2/GDNF KO) as represented by the fold change (FC). Error bars represent the 95% confidence interval. Corrected p-value of control gene expression compared to KO for follistatin and gremlin, 1.2E-3 and 5.5E-3 respectively.

### TGFb dependent ECM remodeling during optic fissure closure

In the next step we addressed the ECM remodeling under the control of TGFb. To this end we more deeply analyzed our microarray data with “DAVID”. We focused on enriched clusters with their terms being in line with ECM and ECM related processes (Fig 6A). We found the terms ECM, secreted proteins, gylcoproteins, biological adhesion and collagen down-regulated (Fig 6A). In the next step we analyzed the expression levels of distinct ECM genes (Fig 6B) like fibronectin1, elastin, epiphycan, keratocan and some collagens. Altogether the analysis indicates a drastic effect TGFb (TGFb/GDNF) has on the ECM composition during the closure of the optic fissure.

**Figure 6:**
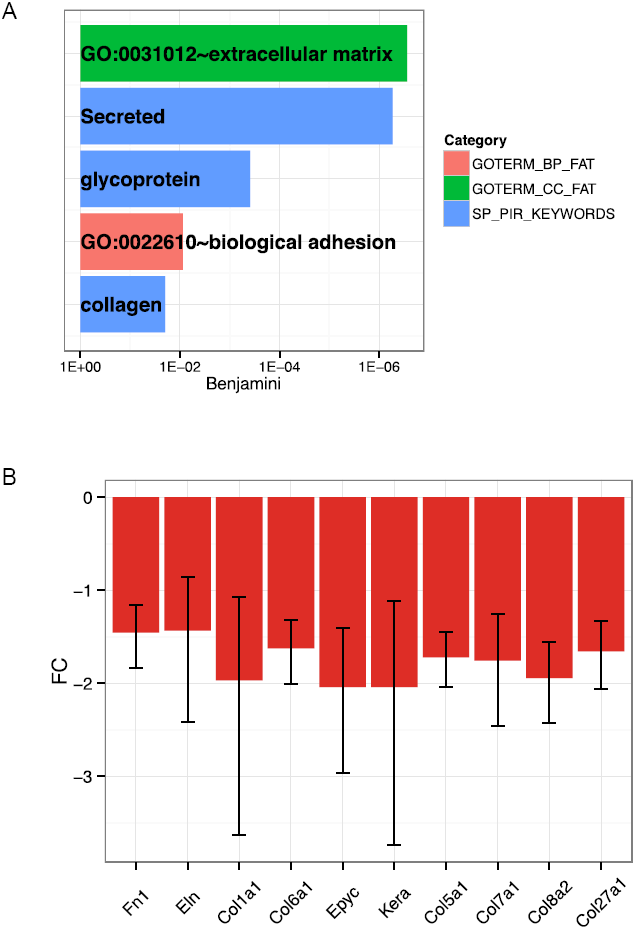
ECM components are the main responsive genes to TGFb2/GDNF KO (A) Selected terms enriched in the set of down-regulated genes in TGFb2/GDNF KO microarray based on the analysis with the tool DAVID. GOTERM_BP_FAT=Biological Process, GOTERM_CC_FAT=Cellular Component, SP_PIR_KEYWORDS=Swiss-Prot Protein Information Resource. (B) Level of misregulation for several ECM related genes represented as fold change (FC). Error bars represent the 95% confidence interval.

## Discussion

At the optic cup stage during vertebrate eye development BMP4 and Vax2 together define dorsal vs. ventral identities of cells in the developing eye (Koshiba-Takeuchi et al., 2000, Sasagawa et al., 2002, Mui et al., 2005, Behesti et al., 2006). A regulated regionalization of the developing eye is important considering the differential retinofugal projections later on. To achieve such a dorso-ventral organization BMP4 is expressed in the dorsal optic cup while Vax2 is expressed in the ventral optic cup.

Recently we described the important role of BMP antagonism during optic vesicle to optic cup transformation (Heermann et al., 2014). BMP signaling must be inhibited for the lens-averted epithelium of the optic vesicle to flow into the developing optic cup and serve as a reservoir for retinal progenitor cells. During this scenario, especially for the ventral part of the optic cup, the forming optic fissure is a key structure (Heermann et al., 2014). Remarkably we found BMP antagonists also expressed at the margins of the optic fissure (this work) suggesting that BMP signaling must be blocked here as well. Our data indicate that TGFb is controlling the expression of the BMP antagonists in this domain. Considering the observations of Zode and colleagues (Zode et al., 2009) who studied the optic nerve head in a glaucoma condition, it is likely that TGFb is rebuilding the ECM at the optic fissure margins, however only if BMP signaling is inhibited in this domain.

Based on our current model (Fig 7) TGFb derived from the lens and the POM induces BMP antagonists in the optic fissure in order to remodel the ECM for enabling the fusion of the meeting optic fissure margins. The lack of TGFb (TGFb2 in mouse) results in a failure of fissure closure, a coloboma (this study).

**Figure 7:**
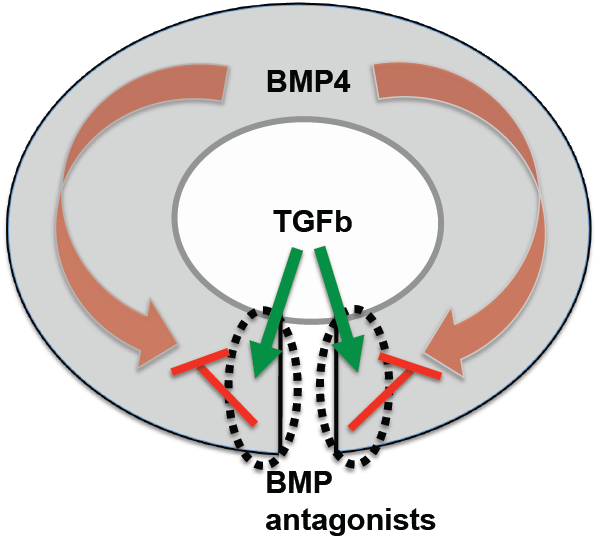
Scheme of proposed signaling network model We propose that TGFb is inducing the expression of BMP antagonists in the optic fissure margins. Blocking BMP signaling in this domain results in ECM remodeling enabling optic fissure fusion.

TGFb was shown to be crucial also for fusion of the palate during development (Sanford et al. 1997, Proetzel et al., 1997). Interestingly TGFb2 and 3 have slightly different functions, since in TGFb2 KO mice a gap is visible (Sanford et al., 1997) and in TGFb3 KO mice the palatal shelves get in touch but do not fuse (Proetzel et al., 1995, Taya et al., 1999). The latter scenario is what we observed while investigating the optic fissure closure in TGFb2 KO embryos. The fissure margins meet, but do not fuse (this study). Interestingly the BMP antagonist Gremlin was recently linked to cleft lips in humans (Al Chawa et al., 2014) indicating that a similar scenario might be active in that context.

Although we started addressing the ECM remodeling during optic fissure closure, it will be a challenge for future studies to further investigate the cell biological basis of the optic fissure closure. This will be including specific changes of ECM components at the fissure margins, which enable fissure fusion. Although the cell biological details are elusive to a great extent, many genes have been linked to coloboma formation (Gregory-Evans et al., 2004, 2013) e.g. Vax2 and Pax2 (Viringipurampeer et al., 2012, Bariberi et al., 2002, Torres et al., 1996). Both genes are expressed in the ventral optic cup. According to our data it is conceivable that the loss of Vax2 and/or Pax2 results in a reduced BMP antagonism, mediating the coloboma in these conditions. However, of course alternatively Vax2 and Pax2 could be more directly linked to the ECM remodeling.

Importantly, in order to apply robustness to the system, BMP antagonists are often expressed redundantly (Khokha et al., 2005, Eggen and Hemmati-Brivanlou 2001, Stottmann et al., 2001) pointing at the importance of their function. According to this it is not surprising that loss of a single BMP antagonist is not resulting in coloboma (Grem1 knock out, not shown).

To study the activity of TGFb signaling in living fish over time we established a TGFb signaling reporter zebrafish line. Next to the identification of the active domain in the optic fissure margins (this study) this line will be very useful to study TGFb signaling during development also of other organ systems or during disease. It will be especially interesting if combined with other pathway reporters (Laux et al., 2011, Schwend et al., 2010).

In summary we showed that TGFb signaling is active in the optic fissure and that loss of a TGFb ligand (TGFb2 in mouse) results in coloboma. Based on our data it is likely that TGFb is inducing BMP antagonists in the fissure margins in order to remodel the ECM and enabling fusion.

## Experimental Procedures

### Transgenic zebrafish

Sequences of Smad binding elements (SBE) in combination with a minimal promoter were cloned into a Gateway 5’ entry vector (Invitrogen). The sequences were kindly provided by Peter tenDijke. A multisite Gateway reaction was performed resulting in an SBE driven GFPcaax construct (SBEsMP::GFPcaax). A Zebrafish line was generated with SB (sleeping beauty) transgenesis according to Kirchmaier et al (Kirchmaier et al., 2013). Shh reporter zebrafish were generated according to Schwend et al (Schwend et al., 2010). The plasmid was kindly provided by Beth Roman.

### Drug treatments

Zebrafish embryos were treated with SB431542 to inhibit TGFb mediated signaling. The substance was dissolved in DMSO.

### Microscopy

Signaling reporter fish were imaged with a Leica SP5 setup, samples mounted in glass bottom dishes (MaTek). For time-lapse imaging embryos were embedded in 1% low melting agarose covered with zebrafish medium an anesthetized with tricain. Left and right eyes were used and oriented to fit the standard views. A stereomircoscope (Olympus/ Nikon) was used for recording brightfiled and fluorescent images of TGFb signaling reporter fish. Whole mount in situ hybridizations were recorded with a stereomicroscope (Olympus) as well as an upright microscope (Zeiss). For time-lapse imaging embryos were embedded in 1% low melting agarose covered with zebrafish medium an anesthetized with tricain. Left and right eyes were used and oriented to fit the standard views.

### Whole mount in situ hybridization

Whole mount in situ hybridizations were performed according to Quiring et al (Quiring et al., 2004). AB/AB zebrafish at approx. 30hpf were used.

### Mice

For this study TGFb2+/- (Sanford et al., 1997) and GDNF+/- (Pichel et al., 1996) mice were used for breeding. Timed matings were performed overnight and the day on which a vaginal plug was visible in the morning was considered as day 0.5. Analyses were restricted to embryonic stages because of perinatal lethality of the individual mutants. For analysis of embryonic tissue, the mother was sacrificed and the embryos were collected by caesarean sectio. All of the experiments were performed in agreement with the local ethical committees.

Genotyping was performed via PCR after gDNA using the following primers:

TGF-b2++/--:
Forward: AAT GTG CAG GAT AAT TGC TGC
Reverse: AAC TCC ATA GAT ATG GGC ATG C
GDNF+/+:
Forward: GAC TAC GGG AGG AGT AGA AG
Reverse: TAT CGT CTC TGC CTT TGT CC
GDNF-/-:
Forward: CCA GAG AAT TCC AGA GGG AAA GGT C
Reverse: CAG ATA CAT CCA CAC CGT TTA GCG G

### Histological analysis

Tissue was processed for paraffin sectioning. Frontal sections of control and TGFb2 - /- (GDNF-/-) were performed and stained with Hämatoxilin and Eosin.

### Microarray data

RNA was extracted from whole eyes of E13.5 embryos (controls and TGFb2-/- (GDNF-/-) respectively). RNA was reverse transcribed, amplified and loaded on Agilent one-color microarray chips. Experiments were performed in triplicates.

Further analysis was performed using R (R Core Team, 2014) and the bioconductor packages Agi4x44PreProcess, limma and mgug4122a.db as annotation database. For background correction we used the followiong parameters: BGmethod = “half”, NORM-method = “quantile”, foreground = “MeanSignal”, background = “BGMedianSignal” and offset = 50. The probes were filtered using the recommended thresholds and afterwards the replicated non-control probes were summarized. Then the method *lmFit* was used to fit a linear model on the arrays. Finally, the differential expression statistics were computed using the methods *eBayes*.

Next only those genes with fold change higher than 1.5 were considered, then a multiple comparison correction was performed on the p-values using the BH (Benjamini & Hochberg) method. The genes with corrected p-value lower than 0.05 were defined as significantly differentially expressed genes.

### DAVID analysis

We used the tool DAVID (Huang et al., 2009a, 2009b) (http://david.abcc.ncifcrf.gov/home.jsp) version 6.7 to find enriched terms on the set of significantly down-regulated genes from the mouse arrays. We provided the Entrez ID of these genes as input and used the default set of databases. For Figure 6 we selected representative terms of the first two clusters found by DAVID.

## Acknowledgements

We want to thank Rolf Zeller for making Gremlin KO mice available as well as for scientific discussion. We also want to thank Gabriela Salinas-Riester and the TAL of the University Göttingen. We also thank Sara Ahlgren for generously making the shh-reporter construct available and Peter tenDijke for the luciferase construct on which our new TGFb reporter zebrafish is based. We thank Lea Schertel for great technical assistance. Finally we want to thank the Deutsche Forschungsgemeinscaft for funding this work.

### Author contributions

SH: conceived the study, carried out experiments, wrote the manuscript, PE: carried out experiments, JM: Microarray data analysis, ER and BR: helped by TGFb KO mouse acquisition, AZ: provided Grem1 KO mice, KK and JW: conceived the study

The authors declare that they do not have a conflict of interest.

